# Effects of age and surface instability on the control of the center of mass

**DOI:** 10.1101/2021.09.08.459390

**Authors:** Maud van den Bogaart, Sjoerd M. Bruijn, Joke Spildooren, Jaap H. van Dieën, Pieter Meyns

## Abstract

During standing, posture can be controlled by accelerating the Center of Mass (CoM) through shifting the center of pressure (CoP) within the base of support by applying ankle moments (“CoP mechanism”), or through the “counter-rotation mechanism”, i.e., changing the angular momentum of segments around the CoM to change the direction of the ground reaction force. Postural control develops over the lifespan; at both the beginning and the end of the lifespan adequate postural control appears more challenging. In this study, we aimed to assess mediolateral balance performance and the related use of the postural control mechanisms in children, older adults and young adults when standing on different (unstable) surfaces. Sixteen pre-pubertal children (6-9y), 17 young adults (18-24y) and eight older adults (65-80y) performed bipedal upright standing trials of 16 seconds on a rigid surface and on three balance boards that could freely move in the frontal plane, varying in height (15-19 cm) of the surface of the board above the point of contact with the floor. Full body kinematics (16 segments, 48 markers, using SIMI 3D-motion analysis system (GmbH) and DeepLabCut and Anipose) were retrieved. Performance related outcome measures, i.e., the number of trials with balance loss and the Root Mean Square (RMS) of the time series of the CoM acceleration, the contributions of the CoP mechanism and the counter-rotation mechanism to the CoM acceleration in the frontal plane and selected kinematic measures, i.e. the orientation of the board and the head and the Mean Power Frequency (MPF) of the balance board orientation and the CoM acceleration were determined. Balance loss only occurred when standing on the highest balance board, twice in one older adult once in one young adult. In children and older adults, the RMS of the CoM accelerations were larger, corresponding to poorer balance performance. Across age groups and conditions, the contribution of the CoP mechanism to the total CoM acceleration was much larger than that of the counter-rotation mechanisms, ranging from 94%-113% vs 23%-38% (with totals higher than 100% indicating opposite effects of both mechanisms). Deviations in head orientation were small compared to deviations in balance board orientation. We hypothesize that the CoP mechanism is dominant, since the counter-rotation mechanism would conflict with stabilizing the orientation of the head in space.

## 1. Introduction

Postural control is the ability to align body segments such that the body center of mass (CoM) stays within safe boundaries above the base of support (BoS) or to bring it back above the BoS after perturbations (Horak, 1987). This is essential for activities of daily living. Postural control depends on the central nervous system to continually integrate and (re)weigh information from visual, vestibular and proprioceptive systems and elicit coordinated muscular responses (Peterka, 2002).

During standing, the base of support is fixed and posture can be controlled by accelerating the CoM by shifting the center of pressure (CoP) within the BoS (Hof, 2007). If CoP shifts are used, the body rotates more or less as a single segment around the ankle(s), often modeled as a single inverted pendulum. Consequently, it has been coined the “CoP mechanism” (Hof, 2007; Horak & Nashner, 1986). Another mechanism to accelerate the CoM, is the “counter-rotation mechanism”, i.e., changing the angular momentum of segments around the CoM to change the direction of the ground reaction force (Hof, 2007). In this mechanism, body segments are rotated with respect to the CoM (Otten, 1999). The hip mechanism, defined by its kinematic characteristics, i.e. the rotation of the trunk and legs around the hip, is one example of the counter-rotation mechanism (Hof, 2007; Otten, 1999). Rotations of other body segments, such as the arms or head can be used in the same way. Therefore, the umbrella term “counter-rotation mechanism” will be used here. Especially when trunk rotations are included, the counter-rotation mechanism may result in head rotations and consequently affect visual and vestibular input (Alizadehsaravi, Bruijn, & van Dieën, 2021). Thus, people may prefer to keep their head orientation stable in space, rather than using rotations involving the head to change angular momentum to control the CoM (Fino, Raffegeau, Parrington, Peterka, & King, 2020).

The use of these postural control mechanisms has been suggested to be direction-specific (Winter, Patla, Prince, Ishac, & Gielo-Perczak, 1998; Winter, Prince, Frank, Powell, & Zabjek, 1996). For anteroposterior control, the CoP can be shifted along the length of the feet by modulating plantar and dorsiflexor muscle activity, while the counter-rotation mechanism in anteroposterior direction could involve hip and trunk flexor/extensor muscle modulation to rotate the trunk and consequently accelerate the CoM or the arms. Accelerating the arms could be very beneficial, as one could potentially accelerate them without having to deaccelerate them. In the mediolateral direction, modulation of evertor and invertor muscle activity allows shifting the CoP along the width of the foot, but in bipedal stance also loading of one, and unloading of the other leg by extensor and flexor muscle respectively, causes a shift of the CoP. The counter-rotation mechanism could involve modulation of hip abductor/adductor and trunk lateroflexor muscle activity to rotate the trunk and accelerate the CoM or swing of the arms. The current work will focus on the use of the CoP and counter-rotation mechanisms in the mediolateral direction, as particularly mediolateral balance impairments have been associated with an increased risk of falling (Maki, Holliday, & Topper, 1994). Falls are the most common cause of childhood injury presented at emergency departments (Sminkey, 2008) and also in older adults, falls are a major cause of injury and even of death (Haagsma et al., 2016; Haagsma et al., 2020). About 30% of community-dwelling Western adults over 65 years of age fall every year (Bath & Morgan, 1999; de Rekeneire et al., 2003; Morrison, Fan, Sen, & Weisenfluh, 2013; Peel, 2011; Stel, Smit, Pluijm, & Lips, 2003). To understand the causes of falling, fundamental knowledge of how healthy ageing affects postural control is of utmost importance.

In healthy young adults, both ankle (invertor/evertor) and hip (abductor/adductor) muscles control mediolateral CoM movement during quiet stance (Day, Steiger, Thompson, & Marsden, 1993; Gatev, Thomas, Kepple, & Hallett, 1999; Winter et al., 1996). Reliance on more proximal muscles has been shown to increase when standing on a compliant or moving support surface (Patel et al., 2008; Riemann, Myers, & Lephart, 2003; van Dieen, van Leeuwen, & Faber, 2015). Standing on such a surface makes proprioceptive information at the ankle less pertinent and changes the effects of ankle moments produced for postural control (Horak & Hlavacka, 2001; MacLellan & Patla, 2006). The mean power frequency (MPF) of CoP displacements is, in young adults, higher when standing on a sway-referenced support surface compared to a rigid surface (Dickin, McClain, Hubble, Doan, & Sessford, 2012). This may suggest that the frequency of postural corrections made using the CoP mechanism is increased in such conditions. In addition, the contribution of the counter-rotation mechanism was shown to be larger when standing on an unstable compared to a rigid surface (van Dieen et al., 2015). As such, it is expected that young adults would rely more on the counter-rotation mechanism with increasing surface instability.

Postural control develops over the lifespan; at both the beginning and the end of the lifespan problems with postural control usually occur. In children, the sensory systems are not completely developed, which diminishes postural control (Steindl, Kunz, Schrott-Fischer, & Scholtz, 2006). Proprioceptive function seems to mature at 3 to 4 years of age and visual and vestibular afferent systems reach adult levels at 15 to 16 years of age (Steindl et al., 2006) or even later (Hirabayashi & Iwasaki, 1995). The integration and reweighting of sensory information do not reach adult levels until the age of 15 (Shams, Vameghi, Shamsipour Dehkordi, Allafan, & Bayati, 2020). During quiet standing, the amplitude of mediolateral CoP displacements and velocities are larger in children between 5 and 7 years old than in older children and adults (Hsu, Kuan, & Young, 2009; Riach & Starkes, 1989). Information on the use of the postural control mechanisms and the effects on sway amplitude and velocity at different surfaces in children is, to the best of our knowledge, missing.

In older adults, sensory and motor impairments, as well as sensory reweighting deficits lead to impaired postural control compared to young adults. Older adults’ ability to adapt to altered sensory conditions such as visual and/or surface perturbations is reduced, and they generally rely more on visual information for postural control than young adults (Bugnariu & Fung, 2007; Teasdale, Stelmach, Breunig, & Meeuwsen, 1991). Consequently, CoP displacement and velocity are larger in older adults (age > 64) compared to young adults both when standing on a rigid surface and when standing on foam (Bergamin et al., 2014; Bugnariu & Fung, 2006). The frequency of movements of the CoP appears similar in older adults compared to young adults (Demura & Kitabayashi, 2006). Older adults appeared to rely more on the counter-rotation mechanism when responding to anteroposterior perturbations than young adults (Wu, 1998). However, in responding to a platform rotation, older adults’ initial lateral trunk and arm movements were directed in the same direction as the platform rotation, suggesting that they anticipated breaking a potential fall, while young adults used arm movements to control CoM movement and recover upright posture (Allum, Carpenter, Honegger, Adkin, & Bloem, 2002). It thus remains an open question whether older adults rely more, or less, on the counter-rotation mechanisms for mediolateral control of standing on rigid and unstable surfaces.

In this study, we assessed mediolateral balance performance and the related use of the postural control mechanisms in children, older adults and young adults. To test if, and how, this changes with ageing, variations in surface instability were used. Understanding the use of the CoP and counter-rotation mechanisms for postural control across the lifespan and when standing on different (unstable) surfaces may have implications for the design of therapeutic interventions that aim to decrease fall incidence.

## 2. Methods

### 2.1. Subjects

Three age groups were compared in this study: pre-pubertal children (6-9 years old), young adults (18-24 years old) and healthy non-falling older adults (65-80 years old). Sample size was calculated for a two-tailed unpaired sample t-test analysis using G*Power (1-β = 0.8, α = 0.05) and an effect size of 1.5 based on previous studies (Masani et al., 2007; Oba, Sasagawa, Yamamoto, & Nakazawa, 2015). The total calculated sample size per group was eight. Potential participants were excluded if they reported any neurological or orthopedic disorder(s), had an uncorrectable visual impairment, were unable to maintain independent and unsupported stance for 60 seconds, had undergone surgery of the lower extremities during the last two years, or took medication that might affect postural control. Additionally, older subjects were excluded if they had experienced two or more falls during normal daily activities in the preceding year or had a cognitive impairment (tested with Mini-Mental state examination (score<24)). Participants gave written informed consent prior to any experimentation. The study protocol was in agreement with the declaration of Helsinki and had been approved by the local ethical committee (CME2018/064, NCT04050774).

### 2.2. Research design

The participants performed bipedal upright standing on a rigid surface and on three balance boards varying in height of the surface of the board above the point of contact with the floor (BB1; 15 cm, BB2; 17 cm and BB3; 19 cm). The balance board was a 48 cm by 48 cm wooden board mounted on a section of a cylinder with a 24 cm radius that could freely move in the frontal plane (Figure 1). The four conditions, were repeated three times in random order, with each trial lasting 16 seconds. For every trial, participants were instructed to stand barefoot on two feet, placed in parallel at hip width and arms along the body. They were asked to look at a marked spot at seven meters distance on the wall in front of them at eye level.

**Figure 1.**
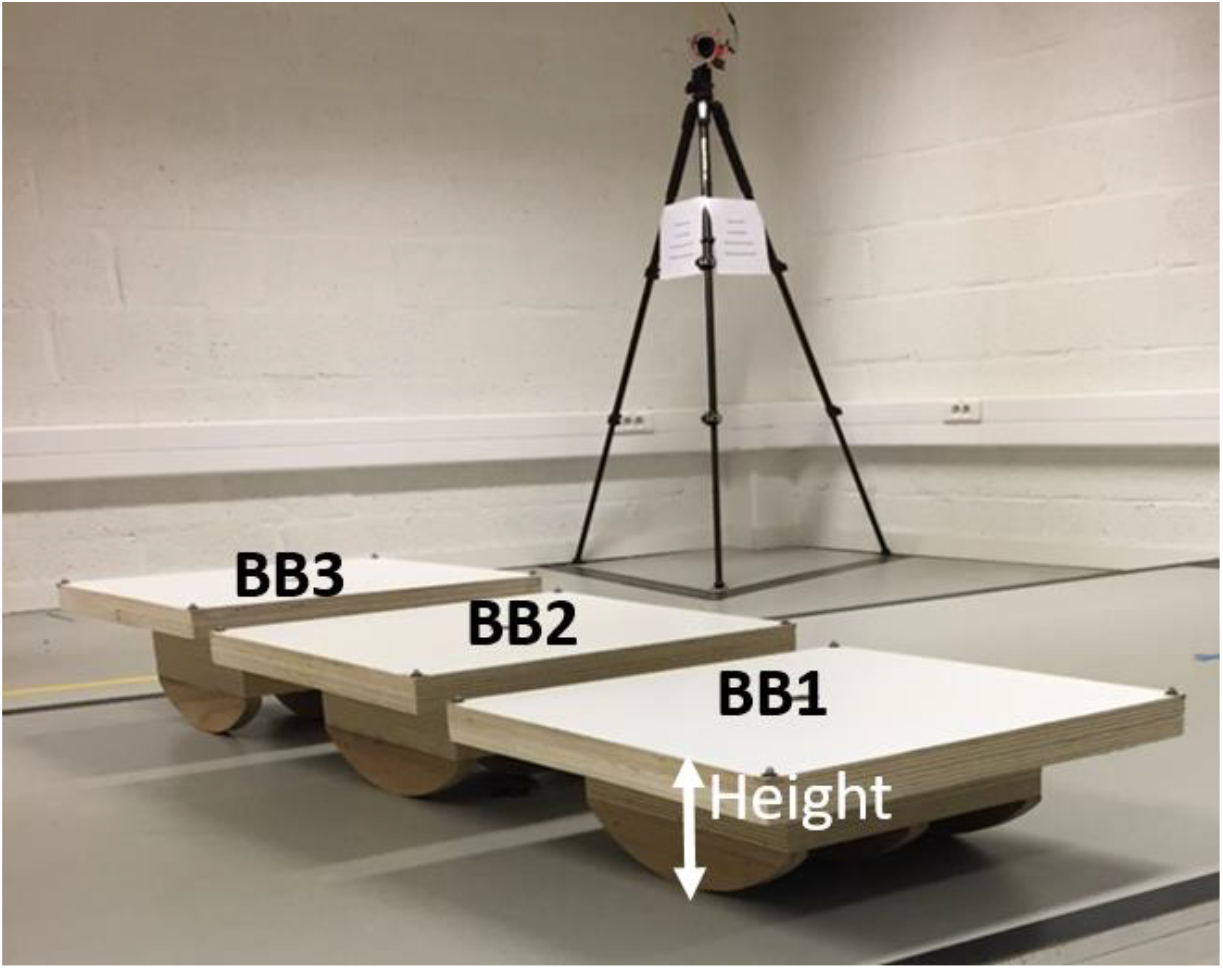
Illustration of the balance boards, which could freely move in the frontal plane, varying in height of the surface of the board above the point of contact with the floor (BB1; 15 cm, BB2; 17 cm and BB3; 19 cm).

### 2.3. Materials and software

A Simi 3D motion analysis system (GmbH) with eight cameras (sample rate:100 samples/sec, resolution: 1152×864 pixels) and 48 retro reflective markers was used (A.1.1. Supplementary materials). Full body 3D kinematics (16 segments) were retrieved using the open-source deep learning python toolboxes DeepLabCut (https://github.com/AlexEMG/DeepLabCut) and Anipose (https://github.com/lambdaloop/anipose) (A.1.1. Supplementary materials). This workflow provided 3D data, which is similar in quality to that obtained with the Simi 3D motion analysis software, yet requires less user interaction (A.1. Supplementary materials).

### 2.4 Data analysis

#### 2.4.1. Performance

A trial was registered as a balance loss if a stepping response or an intervention by a researcher was required in order to maintain stance. The number of balance losses per condition and per age group was recorded as a performance related outcome measure. In case of a balance loss, the trial was excluded from the analysis. Data points were excluded from analysis after participants lifted their foot during a trial or stepped off the balance board, leading to trials lasting shorter than 16 seconds. Next to the number of balance losses, the Root Mean Square (RMS) of the time series of the CoM acceleration in the frontal plane was determined as a measure of performance.

#### 2.4.2. Postural control mechanisms

The contributions of the CoP mechanism and counter-rotation mechanism to the CoM acceleration in the frontal plane, were calculated using Eq. (1), as described by Hof (2007).

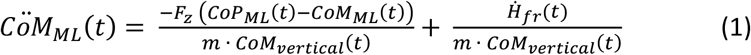

in which *m* is body mass, *CoM*_*ML*_ is the mediolateral (ML) position of the CoM, *CoM*_*vertical*_ is the vertical position of the CoM, *CÖM*_*ML*_ is the double derivative with respect to time of CoM_ML_, *t* is time, *F*_*z*_ is the vertical ground reaction force, *CoP*_*ML*_ is the ML position of the CoP, and 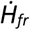 is the change in total body angular momentum in the frontal plane. Here the first part of the right-hand term, 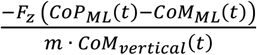, refers to the CoP mechanism and the second part,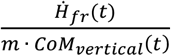, is the ML CoM acceleration induced by the counter-rotation mechanism.

As it was not possible to collect accurate ground reaction forces and CoP when standing on a balance board, the contribution of the CoP mechanism to the *CÖM*_*ML*_ was calculated by substracting, 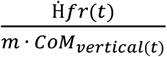, from *CÖM*_*ML*_*(t)*.

The RMS of the time series of the contribution of the CoP mechanism and the counter-rotation mechanism to the *CÖM*_*ML*_ was calculated for each trial. The relative contribution of the CoP mechanism and the counter-rotation mechanism to the *CÖM*_*ML*_ (in %) was calculated by dividing the RMS of each mechanism by *CÖM*_*ML*_, multiplied by 100.

#### 2.4.3. Kinematics

Orientations of the board and head in the frontal plane were calculated. The deviations from the mean orientations were determined by calculating the standard deviation (SD) of the orientation angles. The MPF of the balance board orientation and *CÖM*_*ML*_ were calculated using the pwelch matlab function with a window of 100 data points (1 second), 50% overlap, and nfft of 1000. Before using the pwelch matlab function, the balance board rotation angles were high pass filtered with a second order Butterworth filter with a cutoff frequency of 3 Hz.

### 2.5. Statistics

The results of the three trials of each surface condition were averaged for each participant. Two-way repeated measures ANOVAs were used to determine the effect of Age and Surface as well as their interaction on the RMS and MPF of the *CÖM*_*ML*_, the RMS of the contribution of the CoP and counter-rotation mechanism, the relative contribution of the CoP mechanism and the counter-rotation mechanism to *CÖM*_*ML*_, the SD and MPF of the balance board orientation, and the SD of the head orientation. Post-hoc analyses were performed to determine differences between the different experimental surface conditions (using a Bonferroni correction) and between the different age groups, comparing children and older adults with young adults. Statistical analyses were performed with SPSS(v25) at α<0.05.

## 3. Results

### 3.1. Participants

Sixteen pre-pubertal children between 6-9 years old (10 males, age 8.2±1.1 years old, BMI 15.6±1.5 kg/m^2^), 17 healthy young adults between 18-24 years old (7 males, age 21.9±1.6 years old, BMI 23.5±3.0 kg/m^2^) and eight older adults between 65-80 years old (5 males, age 71.8±4.6 years old, BMI 26.0±3.4 kg/m^2^) participated in this study.

### 3.2. Performance

#### 3.2.1. Balance loss

Balance loss only occurred when standing on the highest balance board, twice in one older adult once in one young adult. No participants had to be excluded because at least one out of the three trials per surface condition per participant was available. During 23 trials (5x BB1, 2x BB2, 6x BB3 and 9x RIGID) some data points were excluded with the result that these trials lasted less than 16 seconds.

#### 3.2.2. Total CoM acceleration

There was a significant interaction between Age and Surface on the RMS of *CÖM*_*ML*_, *F* (6, 114) = 4.423, *p* < 0.001. Post-hoc tests revealed that the RMS of *CÖM*_*ML*_ was significantly smaller during standing on a rigid surface compared to standing on the balance boards in all groups (Figure 2a). Moreover, it was significantly larger in children and older adults compared to young adults during all conditions (Figure 2a), reflective of a poorer balance performance. To further unravel the interaction effect, the effect of age on the difference between the RMS of *CÖM*_*ML*_ during standing on a rigid surface and on BB1, BB2 and BB3 was assessed using independent t-tests. The RMS of *CÖM*_*ML*_ increased significantly more from standing on a rigid surface to standing on BB1, BB2 and BB3 in older adults compared to young adults (*t* (23) = -2.697, *p* = 0.013, *t* (23) = -3.159, *p* = 0.012 and *t* (23) = -2.529, *p* = 0.019, respectively), but not in children compared to young adults (*t* (31) = 0.595, *p* = 0.556, *t* (31) = 0.868, *p* = 0.392 and *t* (31) = 1.878, *p* = 0.072. respectively).

**Figure 2.**
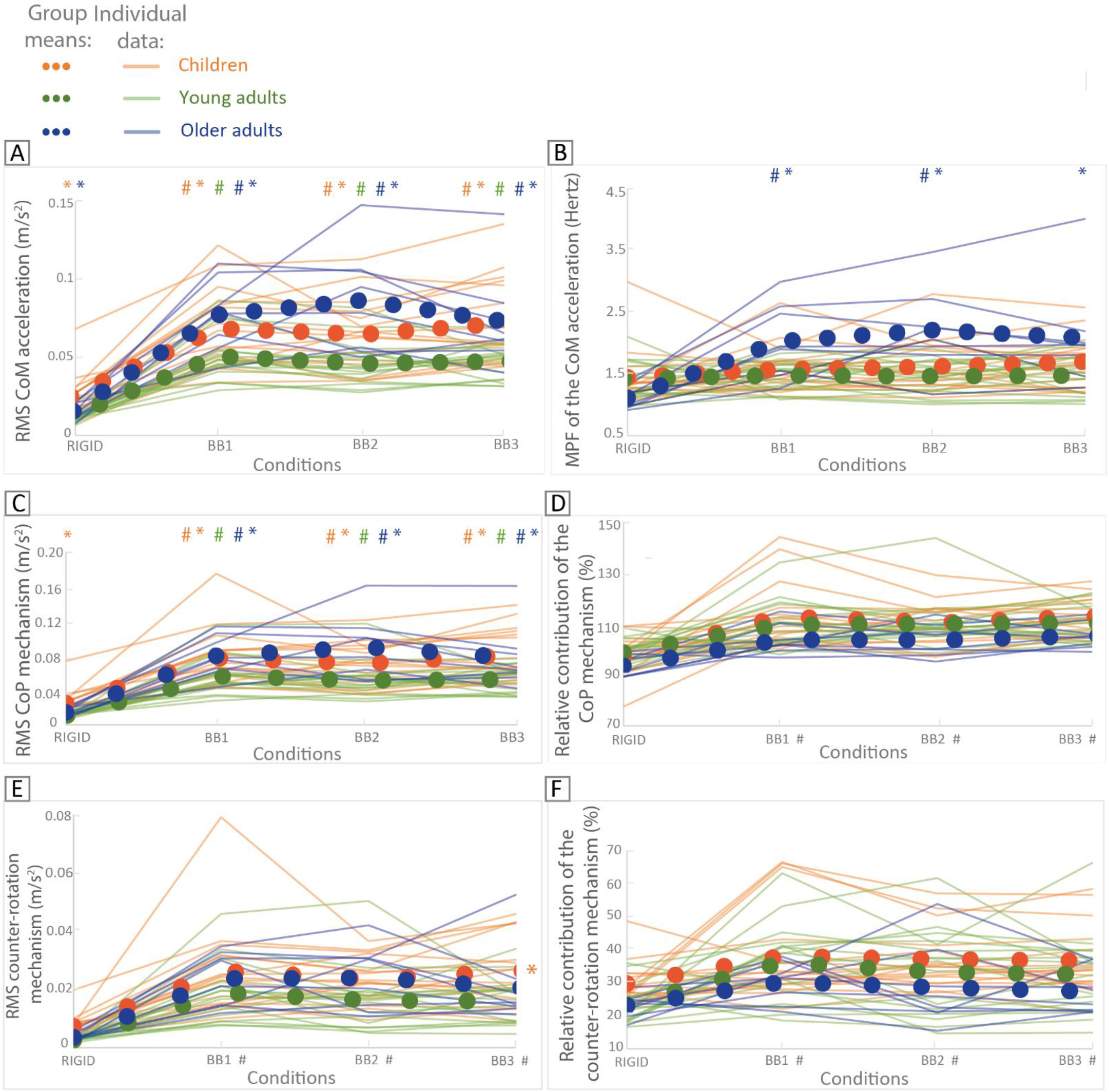
Group means (dotted line) and individual data (solid lines) of the **A)** Root Mean Square (RMS) of the Center of Mass (CoM) acceleration (*CöM*_*ML*_), **B)** Mean Power Frequency (MPF) of *CöM*_*ML*_ (in Hertz (Hz)), **C)** the RMS of the contribution of the CoP mechanism, **D)** the relative contribution of the CoP mechanism to *CöM*_*ML*_, **E)** the RMS of the contribution of the counter-rotation mechanism, **F)** the relative contribution of the counter-rotation mechanism to *CöM*_*ML*_, during standing on a rigid surface (RIGID) and during standing on uniaxial balance boards varying in height (BB1; 15 cm, BB2; 17 cm and BB3; 19 cm) in children (orange), young adults (green) and older adults (blue). # represents significant difference compared to standing on a rigid surface. * represents significant difference compared to young adults, with the group tested identified by the color code.

#### 3.3. Postural control mechanisms

There was a significant interaction between Age and Surface on the RMS of the contribution of the CoP mechanism, *F* (6, 114) = 2.822, *p* = 0.013. Post-hoc tests revealed that the contribution of the CoP mechanism was significantly smaller during standing on a rigid surface compared to standing on the balance boards in all groups (Figure 2c). Moreover, it was significantly larger in children and older adults compared to young adults during all balance board conditions. When standing on a rigid surface, the contribution of the CoP mechanism was significantly larger in children compared to young adults (Figure 2c). There was no significant difference between young adults and older adults when standing on a rigid surface (*p* = 0.088). To further unravel the interaction effect, the effect of Age on the difference between the RMS of the contribution of the CoP mechanism during standing on a rigid surface and standing on BB1, BB2 and BB3 was assessed using independent t-tests. The increase in the RMS of the contribution of the CoP mechanism from standing on a rigid surface to BB3 was significantly larger for children compared to young adults (*t* (31) = 2.153, *p* = 0.039), but the increase from standing on a rigid surface to BB1 and BB2 was not significant different between children and young adults (*t* (31) = 0.78, *p* = 0.438 and *t* (31) = 0.871, *p* = 0.391). The increase in the RMS of the contribution of the CoP mechanism from standing on a rigid surface to BB2 and BB3 was significantly larger in older adults compared to young adults (*t* (23) = -3.105, *p* = 0.005 and *t* (23) = -2.128, *p* = 0.044 respectively), but the increase from standing on a rigid surface to BB1 was not significant different between older adults and young adults (*t* (23) = -2.027, *p* = 0.054.

The relative contribution of the CoP mechanism to *CÖM*_*ML*_ ranged from 94%-113%. (Figure 2d). The average relative contribution increased from standing on a rigid surface to standing on the balance boards (effect of Surface, *F* (2.010, 76.370) = 32.621, *p* = < 0.001, Figure 2d). There was no significant interaction between Age and Surface and the effect of Age was not significant.

The RMS of the contribution of the counter-rotation mechanism was significantly smaller when standing on a rigid surface than when standing on the balance boards (effect of Surface, *F* (3, 114) = 53.863, *p* < 0.001). Moreover, the contribution of the counter-rotation mechanism was significantly larger in children compared to young adults (effect of Age, F (2, 38) = 4.076, p = 0.025, Figure 2e). There was no significant interaction between Age and Surface.

The relative contribution of the counter-rotation mechanism to *CÖM*_*ML*_ ranged from 23%-38%. (Figure 2f). The relative contribution of the counter-rotation mechanism increased from standing on a rigid surface to standing on the balance boards (effect of Surface, *F* (1.993, 75.727) = 10.813, *p* = < 0.001, Figure 2f). There was no significant interaction between Age and Surface and the effect of Age was not significant.

### 3.4 Kinematics

#### 3.4.1. Balance board orientation

The SD of balance board orientation was larger in children and older adults compared to younger adults (effect of Age, *F* (2, 38) = 8.256, *p* = 0.001, Figure 3b), implying larger deviations from the mean board orientation. There was no significant interaction between Age and Surface and the effect of Surface was not significant.

**Figure 3.**
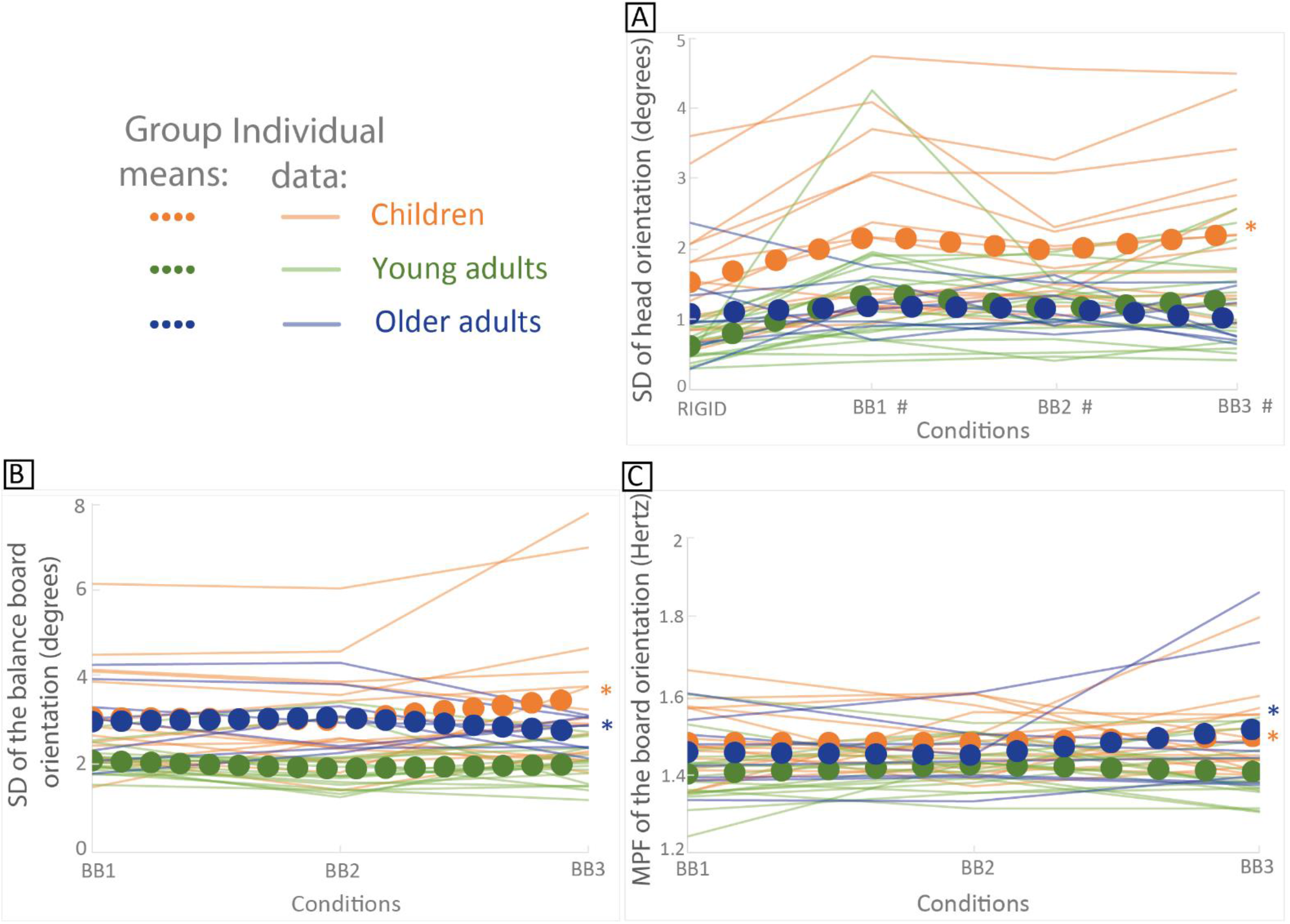
Group means (dotted lines) and individual data (solid lines) of the **A)** standard deviation (SD) of the head orientation (in degrees), **B)** SD of the balance board orientation (in degrees), **C)** Mean Power Frequency (MPF) of the board orientation (in Hertz (Hz)), during standing on uniaxial balance boards varying in height (BB1; 15 cm, BB2; 17 cm and BB3; 19 cm) in children (orange), young adults (green) and older adults (blue). * represents significant difference compared to young adults, with the group tested identified by the color code. # represents significant difference compared to standing on a rigid surface.

#### 3.4.2. Head orientation

The SD of head orientation was significantly smaller when standing on a rigid surface than when standing on the balance boards (effect of Surface, *F* (2.521, 95.805) = 10.524, *p* < 0.001, Figure 3a). Moreover, the SD of head orientation was significantly larger in children compared to young adults, while there was no difference between older and younger adults (effect of Age, F (2, 38) = 7.329, p = 0.002, Figure 3a). There was no significant interaction between Age and Surface.

#### 3.4.3. Frequency of CoM accelerations and balance board rotations

Age and Surface had a significant interaction effect on the MPF of *CÖM*_*ML*_, *F* (4.417, 83.927) = 7.316, *p* < 0.001. In older adults, the MPF of *CÖM*_*ML*_ was significantly lower during standing on the rigid surface compared to standing on BB1 and BB2 (Figure 2b). The MPF of *CÖM*_*ML*_ was significantly higher in older adults compared to young adults during all balance board conditions (Figure 2b). There was no difference between children and younger adults.

The MPF of balance board orientation was larger in children and older adults compared to younger adults (effect of Age, *F* (2,38) = 5.081, *p* = 0.011, Figure 3c), implying increased frequency of the balance board rotations. There was no significant interaction between Age and Surface and the effect of Surface was not significant.

## 4. Discussion

The aim of this study was to obtain insight in differences in balance performance and the related use of postural control mechanisms between children, older adults and young adults during standing on a rigid surface and on balance boards which could freely move in the frontal plane. We found an increased RMS of CoM accelerations both in children and older adults compared to young adults, implying worse balance performance. Additionally, we found that the relative contribution of the CoP mechanism to the total CoM acceleration was around 100% in all surface conditions and age groups

### 4.1. Effects of surface instability

Balance loss only occurred when standing on the highest balance board. The RMS of CoM accelerations was larger during standing on a balance board than on the rigid surface. These performance outcome measures, as well as increased RMS-values and contribution of the CoP and counter-rotation mechanism when standing on the balance boards reflect that standing on the balance board was indeed more challenging than standing on the floor (van Dieen et al., 2015). This can be explained by the decrease of pertinent proprioceptive information from ankle muscles and a reduction of the effectiveness of ankle moments for postural control when standing on a balance board (Horak & Hlavacka, 2001). Furthermore, participants rotated their head more when standing on the balance boards compared to standing on a rigid surface, probably due the moving support surface. However, the standard deviation of the head rotation was limited to only 2.2 degrees.

Surprisingly, the increasing height of the balance boards did not cause a further deterioration of balance performance, change in use of postural control mechanisms or differences in kinematics. This was unexpected, as the stability of a person on a balance board and therefore the difficulty of the task is affected by the height of the surface of the board above the point of contact with the floor (B.1. Supplementary materials, Eq. (B.1.)). Most likely, the differences between the height of the balance boards were not big enough to cause differences. Overall, the standard deviation of the board orientation ranged between 1.9 and 3.6 degrees, suggesting that the challenge imposed was limited for all participants.

### 4.2. Age effects

#### 4.2.1. Performance and kinematics

One older adult lost balance twice and one young adult once when standing on the highest balance board. The limited number of balance losses when standing on the highest balance board and the absence of balance losses on the other balance boards suggests that all participants had good capacity to keep standing on the balance boards. In children and older adults, sensorimotor control is worse compared to young adults, hence an increased RMS of CoM accelerations, corresponding to a deterioration of balance performance, and larger deviations from the mean board orientation compared to young adults were expected (Bugnariu & Fung, 2006; Hirabayashi & Iwasaki, 1995; Shams et al., 2020; Steindl et al., 2006; Teasdale et al., 1991). In older adults, the increase in RMS of CoM accelerations from standing on a rigid surface to standing on a balance board was significantly larger compared to young adults. Thus, the effect of aging on balance performance was amplified by the task difficulty, suggesting that a decrease of pertinent proprioceptive information and/or a reduction of effectiveness of ankle moments for postural control poses more of a challenge for the older adults. The differences between older and young adults during standing on a rigid surface or a balance board are in line with previous studies (Bergamin et al., 2014; Sturnieks, Arnold, & Lord, 2011). Moreover, the differences in postural control between children and young adults were in line with previous literature using foam as an unstable surface (Hsu et al., 2009; Riach & Starkes, 1989).

The difference in RMS of CoM accelerations between children and young could also be explained by significantly more head rotation in children in addition to the reduction in sensorimotor control. More head rotation could be self-generated and form a sign of less attention in children, which is common in children compared to young adults (Huang-Pollock, Carr, & Nigg, 2002; Wickens, 1974). Self-generated head rotation may be disadvantageous, as it leads to changing visual and vestibular inputs, requiring more processing to discern between body motion and self-imposed head motion (Khan & Chang, 2013). However, the difference in head rotation between children and young adults was less than 1 degree.

The standard deviation and the MPF of the board orientation were significantly higher in children compared to young adults. This indicates more frequent movements of the board over larger angles. This could be due to faster feedback, due to shorter neural pathways in children. However, when standing on the balance boards, the MPF of the CoM accelerations was not significantly larger in children compared to young adults. Thus, the increased frequency of balance board rotations, potentially reflecting an increased frequency of control actions, did not result in an increased frequency of CoM corrections, or improved balance performance. In older adults, the RMS of CoM accelerations and standard deviation of the board orientation were higher compared to young adults, as were the MPF of the CoM accelerations and of the board orientation. This implies more frequent movements of the balance board over larger angles, decreased balance performance and an increased frequency of CoM corrections in older adults compared to young adults. It should be kept in mind that board rotations can reflect corrective actions as well as perturbations due neuromuscular noise. Overall, our results suggest that both children and older adults had more difficulty in controlling the board as compared to young adults.

#### 4.2.2. Postural control mechanisms

The relative contribution of the CoP mechanism to the total CoM acceleration was around 100% (ranging from 94%-113%) and the relative contribution of the counter-rotation mechanism was around 30% (ranging from 23%-38%). Overall, the contribution of the counter-rotation mechanism was limited and not always in the same direction as the contribution of the CoP mechanism, as the RMS values of the contribution of the CoP mechanism were often larger than the RMS values of the total mediolateral CoM acceleration and the relative contributions of the CoP mechanism were around 100%. However, the desired direction is unclear. It could be that segmental rotations were used to achieve a proper orientation of segments such as regulating the orientation of the head in space, rather than controlling the CoM acceleration/position (Alizadehsaravi et al., 2021). Overall, the participants, even the children, kept their head quite stable, in comparison to the board. This suggests that people prefer to keep the head stable, to maintain a constant visual and vestibular input, rather than using upper body rotations as a counter-rotation mechanism to control the CoM. Larger contributions of counter-rotation were found in unipedal stance on a balance board, but this was to a large part generated by the free leg (van Dieen et al., 2015). A limited use of the counter-rotation mechanism to control the CoM was also found in our previous study, in which we found that using the counter-rotation mechanism would actually have interfered with the gait pattern (van den Bogaart, Bruijn, van Dieen, & Meyns, 2020). Combined, these results suggest that the contribution of the counter-rotation mechanism to control the CoM is limited.

This may be because rotational accelerations of body parts may interfere with other task constraints and because they need to be reversed leading to the opposite effect on the acceleration of the body center of mass.

### 4.3. Limitations

Trials leading to balance loss were excluded, which implies that we somewhat overestimated the performance in one older adult and one young adult. However, only three times balance losses occurred and no participants had to be excluded, because at least one out of the three trials per condition per participant was available. Additionally, it should be considered that differences in body mass and height of the center of mass influence the difficulty of standing on the balance boards (B.1.1. Supplementary materials). These could be considered confounding variables, especially when comparing the children to the young adults, but on the other hand this is part and parcel of the age difference. Finally, it was not possible to collect accurate ground reaction forces and CoP when standing on a balance board. However, the contribution of the CoP mechanism could be calculated based on the other terms in Eq. (1).

## 5. Conclusion

In children and older adults, the RMS of CoM accelerations and deviations from the mean balance board orientation were larger, corresponding to poorer balance performance, probably related to less optimal sensorimotor control compared to young adults. The contribution of the CoP mechanism to the total CoM acceleration was around 100%, regardless of age and condition, while the contribution of the counter-rotation was only around 30%. In addition, across groups and conditions, deviations in head orientation were smaller than deviations in balance board orientation. Our results suggest that the CoP mechanism is dominant, likely because the counter-rotation mechanism would conflict with stabilizing the orientation of the head in space.

## 6. Acknowledgements

The authors gratefully acknowledge The Expertise centre for Digital Media (EDM) for technical support and Kimmy Daenen and Charlotte Uwents for their help during the experiments.

## A.1. Supplementary materials

### A.1.1. Workflow DeepLabCut and Anipose

First, the DeepLabCut toolbox was used to extract minimally 115 frames per camera from multiple videos of different persons varying in gender and age (Table A.1., Figure A.1.) (Mathis et al., 2018; Nath et al., 2019). A full-body human model consisting of 48 markers was manually labelled in the extracted frames (Figure A.1.). These data were used to train a deep neural network (resnet_50) in DeepLabCut for each camera with a training fraction of 0.95 and the following learning rates; lr=0.005 for 10k, then lr=0.02 for 430k, lr=0.002 for 730k and lr=0.001 for 1.3M iterations (Table A.1.). The trained networks were used to track 2D coordinates from the videos of the eight cameras (Figure A.1.). To acquire 3D coordinates, Anipose was used to triangulate with optimization across camera views using the calibration data as extracted from the standard Simi calibration (Figure A.1.). A Simi motion wand and L-frame calibration was performed to calibrate the cameras and the calibration file was converted into the “openCV” format which Anipose uses, to be able to triangulate across the camera views (https://osf.io/7hn8z/). A note regarding this conversion is, that is was not completely possible to convert the Simi distortion parameters to open CV format because the distortion values retrieved via the Simi calibration are according a Simi-internal format.

The configuration file parameters for triangulation of the data in Anipose can be found at https://osf.io/7hn8z/.

**Table A.1.** Number of labelled frames, trained iterations and test errors per camera.

|  | Labelled frames | Iterations | Test error (in pixels, with p-cutoff of 0.8) |
| --- | --- | --- | --- |
| *Cam 1* | 125 | 550k | 2.16 |
| *Cam 2* | 245 | 450k | 2.34 |
| *Cam 3* | 155 | 650k | 2.05 |
| *Cam 4* | 115 | 450k | 2.52 |
| *Cam 5* | 165 | 450k | 2.15 |
| *Cam 6* | 210 | 450k | 2.3 |
| *Cam 7* | 269 | 500k | 2.39 |
| *Cam 8* | 160 | 500k | 2.46 |

For training and analyzing the videos with DeepLabCut a computer with a MSI GeForce GTX 1080 Ti graphics card (8GB) was used.

**Figure A.1.**
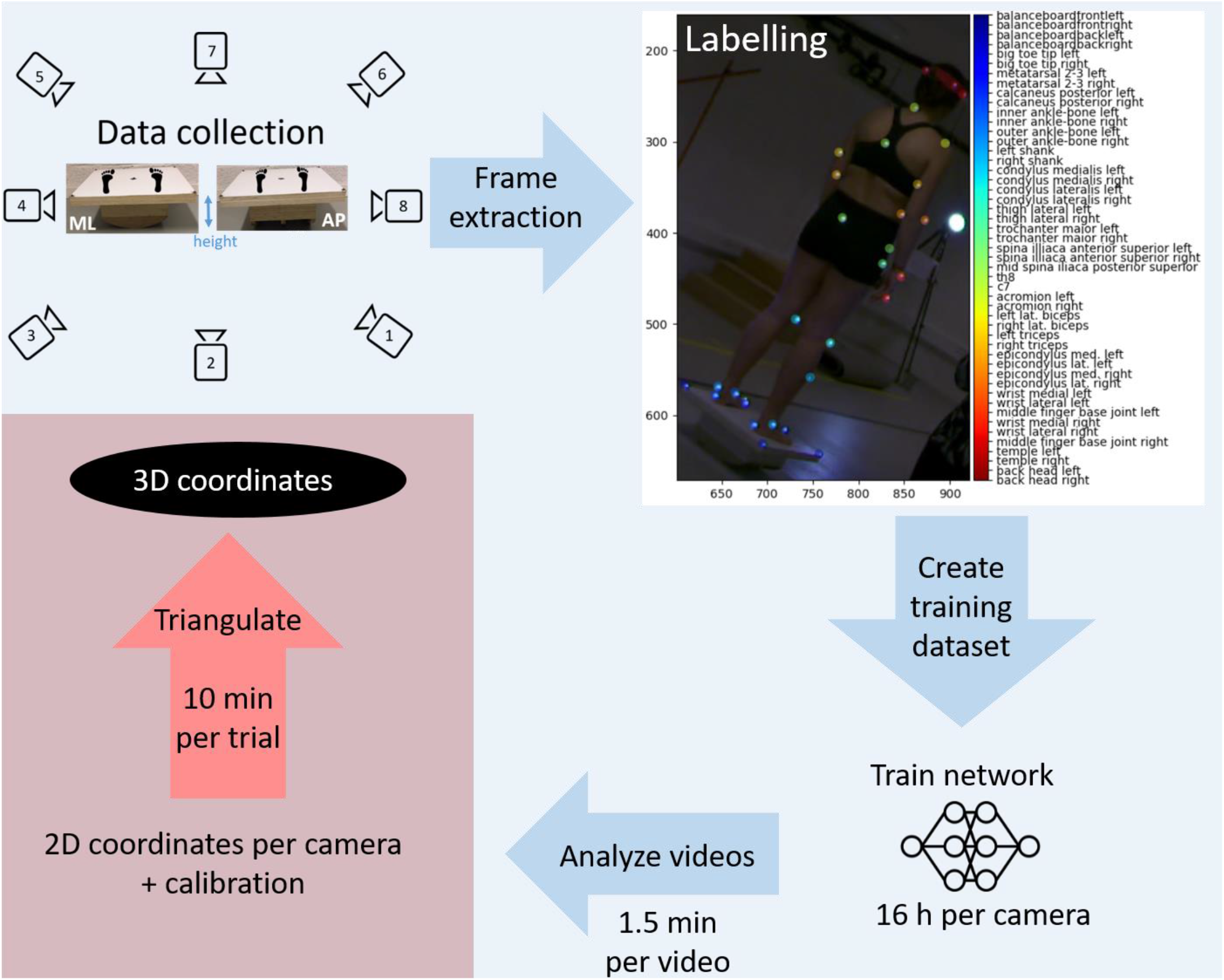
Illustration of the workflow using deep learning (DeepLabCut and Anipose) to retrieve full-body 3D coordinates. DeepLabCut was used for the blue framed steps and Anipose was used for the red framed step in this workflow.

### A.1.2. Comparison Deeplabcut-Anipose and Simi 3D Motion analysis software

Next to retrieving the 3D coordinates using deep learning (Anipose and DeepLabCut), 3D coordinates using Simi 3D Motion analysis software were retrieved of trials of standing on a uniaxial balance board, creating an unstable support in the frontal plane, with a height of 19 cm of 4 young adults, 3 children and 4 older adults. The retrieved 3D coordinates via Simi were low-pass filtered at 10 Hz using a 2^nd^ order Butterworth filter.

From the 3D data of both systems, we created a 3d model, and calculated full body Center of Mass (CoM) displacement, velocity and acceleration (https://osf.io/7hn8z/). The 3D Root Mean Squares (RMS) of the CoM displacement, velocity and acceleration were calculated (Table A.2.). In figure A.2. an example is displayed of a comparison between time series of the CoM displacement, velocity and accelerations with corresponding RMS determined via deep learning (Anipose-Deeplabcut) and Simi 3D motion analysis software.

**Table A.2.** Comparison of deep learning (Anipose-DeepLabCut) and Simi software based calculations of 3D Root Mean Squares (RMS) of Center of Mass (CoM) displacements (in cm), velocities (in cm/s) and accelerations (in cm/s^2^). With x corresponding to mediolateral movement, y corresponding to anteroposterior movement and z corresponding to vertical movement.

| | Displacement (cm) | | | Velocity (cm/s) | | | Acceleration ($\text{cm/s}^2$) | | |
| --- | --- | --- | --- | --- | --- | --- | --- | --- | --- |
|  | x | y | z | x | y | z | x | y | z |
| **mean RMS deep learning** | 45.00 | 26.24 | 107.40 | 1.30 | 0.96 | 0.36 | 5.41 | 5.60 | 3.87 |
| **mean RMS Simi** | 45.19 | 26.14 | 107.28 | 1.29 | 0.90 | 0.36 | 5.47 | 5.95 | 4.45 |
| **Absolute RMS difference between deep learning and Simi based calculations** | 0.19 | 0.09 | 0.12 | 0.01 | 0.06 | 0.00 | 0.06 | 0.35 | 0.58 |

**Figure A.2.**
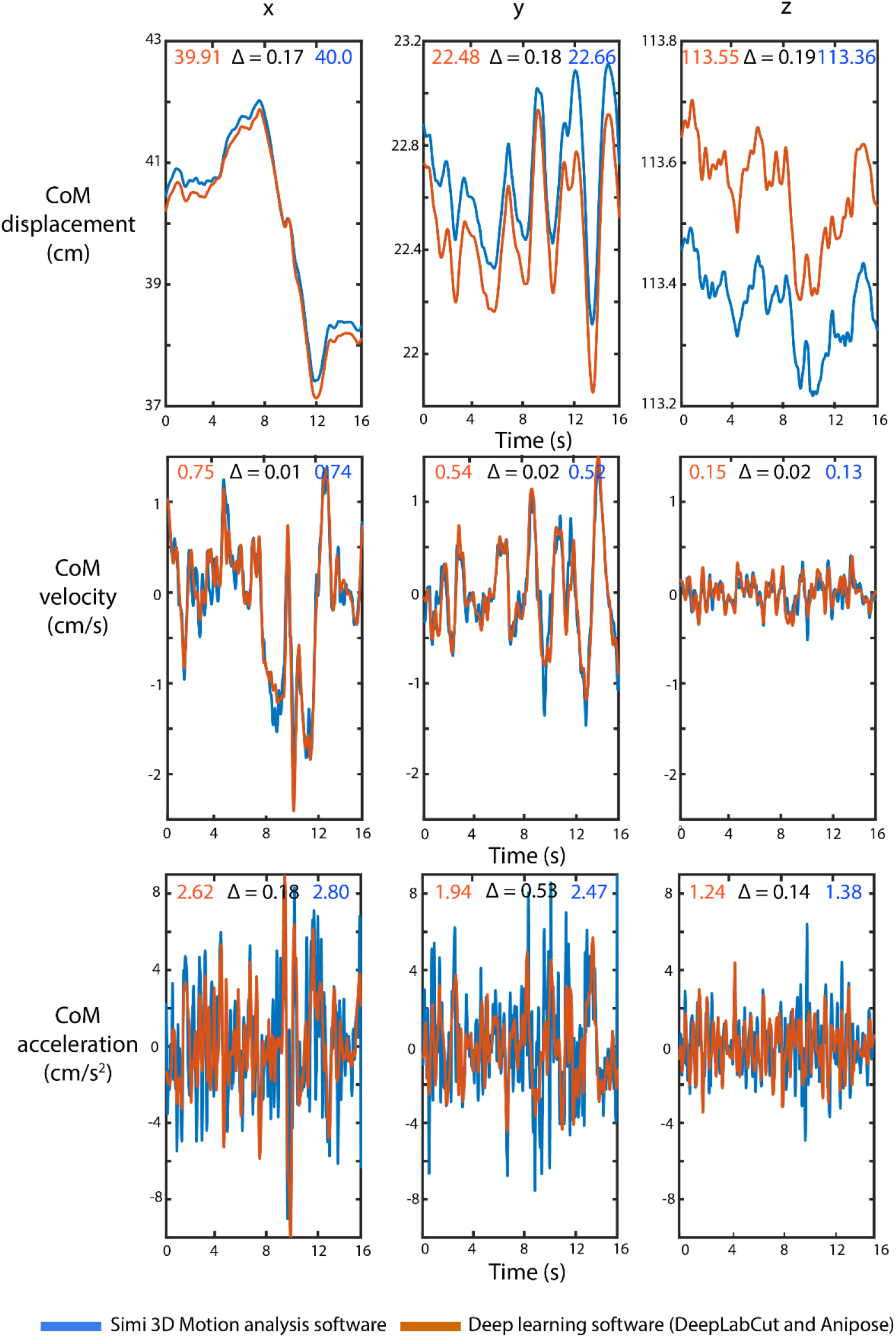
Comparison of Center of Mass (CoM) displacement, velocity and acceleration between calculation based on 3D coordinates via Simi 3D Motion analysis software (in blue) and deep learning software (DeepLabCut and Anipose, in red). The Root Mean Squares (RMS) for Simi 3D Motion analysis software are displayed in red text and for deep learning software in blue text. The differences between the RMSs of the two software’s are displayed with delta (Δ) in black text.). With x corresponding to mediolateral movement, y corresponding to anteroposterior movement and z corresponding to vertical movement.

## B.1. Supplementary materials

### B.1.1. Illustration of the determinants of the difficulty of a balancing task on a balance board with a cylindrical or spherical contact surface

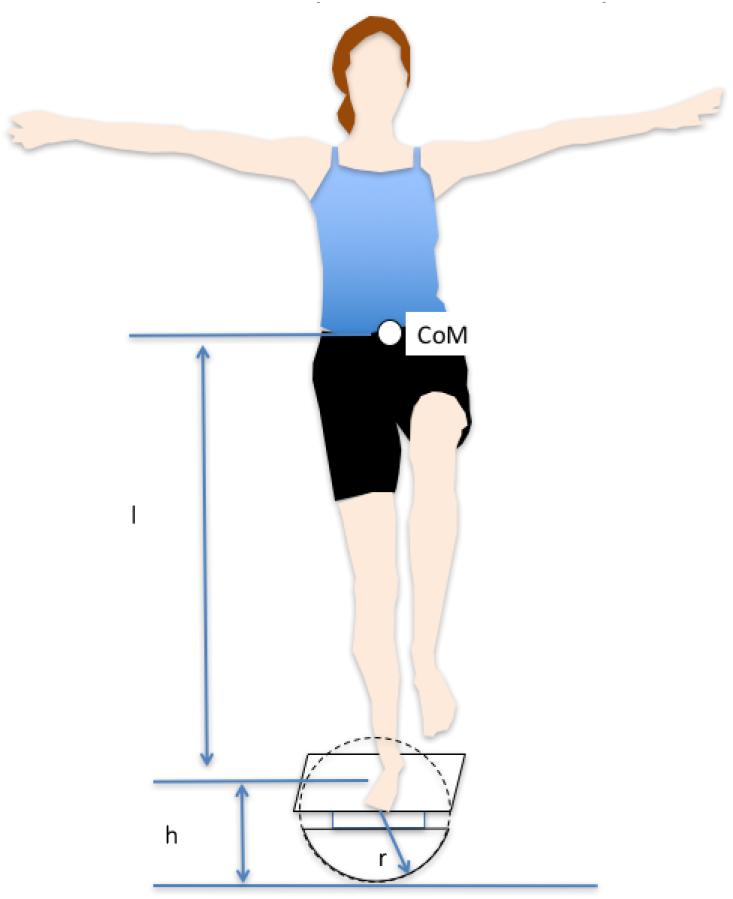

#### Stability of the complete system dM/dϕ

With a rotation of the board over an angle ϕ, the center of mass (CoM) will be displaced horizontally over a distance sin(ϕ)(h+l). This causes a destabilizing moment of gravity that for small angle varies with the angle as: dM/dϕ = mg(h+l). At the same time, the point of support will shift with a change in board angle ϕ as rϕ. Ignoring horizontal forces this leads to an opposite moment that varies with board angle as: dM/dϕ = -mgr. The system is critically stable when r=h+l (as it would be for a completely spherical object). This shows that difficulty of the task can be expressed as:

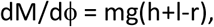

which will be equally affected by increases in h or decreases in r of equal magnitude.

#### Controllability through ankle angle da/dϕ or ankle moment da/dM_a_

Given Eq. (B.1.)

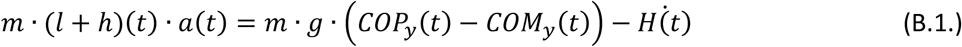

Taking CoM_y_ = 0 as reference and ignoring angular momentum changes, this simplifies to:

a = gCoP/(l+h)= g rϕ /(l+h), which yields:

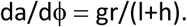

If inertia of balance board is constant, ankle moment M_a_ is related to dϕ, as dϕ = dM_a_dt^2^/I_board_.

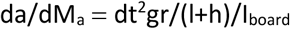

So controllability shows stronger dependence on r than on h, if h is not much larger than l.

